# High prevalence of antibiotic resistance in *Helicobacter pylori* isolates from Iran: importance of functional and mutational analysis of resistance genes and virulence genotyping

**DOI:** 10.1101/569814

**Authors:** Nastaran Farzi, Abbas Yadegar, Hamid Asadzadeh Aghdaei, Amir Sadeghi, Mohammad Reza Zali

## Abstract

The high prevalence of antibiotic resistance in *Helicobacter pylori* has become a great challenge in Iran. The genetic mutations that contribute to the resistance have yet to be precisely identified. This study aimed to investigate the prevalence of antibiotic resistance and virulence markers in Iranian *H. pylori* isolates and to analyze if there is any association between resistance and genotype. Antibiotic susceptibility patterns of 33 *H. pylori* isolates were investigated against metronidazole, clarithromycin, amoxicillin, rifampicin, ciprofloxacin, levofloxacin and tetracycline by the agar dilution method. The *frxA, rdxA, gyrA, gyrB* and 23S rRNA genes of the isolates were sequenced. The virulence genotypes were also determined using PCR. Metronidazole resistance was present in 81.8% of the isolates, followed by clarithromycin (36.4%), ciprofloxacin (36.4%), amoxicillin (30.3%), rifampicin (30.3%), levofloxacin (27.3%) and tetracycline (6.1%). Most of the metronidazole-resistant isolates carried frameshift mutations in both *frxA* and *rdxA* genes, and premature termination was occurred in positions Q5Stop and Q50Stop, respectively. Amino acid substitutions M191I, G208E, and V199A were predominantly found in *gyrA* gene of fluoroquinolone-resistant isolates. A2143G and C2195T mutations of 23S rRNA were found in four isolates. Interestingly, significant associations were demonstrated between intact *cag*PAI and resistance to rifampicin (*P* = 0.027), and between susceptibility to amoxicillin and *cag*PAI intactness (*P* = 0.016). The prevalence of *H. pylori* antibiotic resistance is high in our region, particularly that of metronidazole, clarithromycin, ciprofloxacin and multidrug resistance. Occurrence of mutations in resistance genes were involved in the development of resistance, especially in less virulent isolates.

## Introduction

*Helicobacter pylori* (*H. pylori*) is known as the most common human pathogen infecting more than half of the world’s population.^1,2^ Early eradication based therapies have been proven to regress the *H. pylori*-associated diseases.^3,4^ However, the efficacy of eradication treatments has been extremely compromised primarily due to increased resistance to antimicrobial agents in many countries.^5-8^

Today, first-line standard triple therapy is the most widely used eradication treatment for *H. pylori* infection, which typically comprises two of three antibiotics including amoxicillin, clarithromycin, and metronidazole in combination with one proton pump inhibitor (PPI).^3,9^ However, the use of levofloxacin or ciprofloxacin in fluoroquinolone containing triple therapy and bismuth-based quadruple therapy have also been suggested as second-line therapies after the failure of the clarithromycin-containing regimens.^10-12^ Furthermore, tetracycline and rifampicin are among the common antibiotics that have been used in several rescue therapies recommended in eradication of *H. pylori* infection.^13-15^

Previous studies have demonstrated that numerous point mutations resulting from genetic plasticity within the chromosomal genes, is the main antibiotic resistance mechanism among *H. pylori* strains in various geographic regions.^5,6,16-18^ Primary resistance to clarithromycin has been mainly associated with point mutations in the peptidyl transferase region encoded in domain V of 23S rRNA. Most of these mutations include nucleotide substitutions involving an adenine to guanine transition at positions 2142 and 2143, and to a lesser extent an adenine to cytosine transversion at position 2142.^8,10,19^ However, several other mutations associated with clarithromycin resistant isolates seem to be emerging.^20,21^ The mechanisms of metronidazole resistance in *H. pylori* are frequently attributed to inactivating mutations in *rdxA* and *frxA* genes.^22,23^ On the other hand, mutational changes leading to various amino acid substitutions that confer fluoroquinolone resistance have been located in different positions of quinolone-resistant determining region (QRDR) of *gyrA* and *gyrB* genes.^19,24^

Apart from aforementioned mechanisms of resistance developed by *H. pylori* strains to the major antibiotics used in the treatment of infection, other factors such as the virulence genotype status of bacteria have been reported to affect drug resistance.^25-28^ However, the exact underlying mechanisms involved in the crosstalk of *H. pylori* virulence and antimicrobial resistance remained to be clarified.

Hence, the focus of the present study was to evaluate the antibiotic susceptibility patterns and underlying resistance mechanisms of *H. pylori* strains isolated from Iranian patients with different gastric diseases. Furthermore, we determined the presence of genetic mutations that are associated with antibiotic resistance. We also examined the possible association between resistance profiles and a panel of virulence genotypes.

## Methods

### Patients and *H. pylori* isolates

Antral biopsies were collected for culture from 78 patients who underwent upper gastroduodenal endoscopy at Taleghani Hospital in Tehran from February 2016 to September 2016. Patients were excluded if they were taking eradication therapy for *H. pylori*, PPIs or H_2_-receptor blockers, and any antibiotics used for other infections within two weeks prior to enrolment. The study protocol was approved by the Ethical Review Committee of the Research Institute for Gastroenterology and Liver Diseases at Shahid Beheshti University of Medical Sciences (Project No. IR.SBMU.RIGLD.REC.1395.878). All experiments were performed in accordance with relevant guidelines and regulations recommended by the institution and informed consents were obtained from all subjects and/or their legal guardians prior to sample collection.

The biopsy specimens were smeared on Brucella agar (Merck, Germany) plates containing 7% horse blood (v/v), 10% fetal calf serum (FCS), Campylobacter-selective supplement and amphotericin B (2.5 mg/l). The inoculated plates were incubated at 37°C in a CO_2_ incubator under microaerophilic atmosphere containing approximately 5% O_2_, 10% CO_2_ and 85% N_2_ for 3-10 days. The *H. pylori* was identified by colony and microscopic morphology, positive catalase, oxidase, and urease tests and confirmed by molecular assays.^29,30^

### Antibiotic susceptibility testing

The antibiotic susceptibility of the *H. pylori* strains was assessed by the agar dilution method against a panel of 7 antibiotics purchased from Sigma-Aldrich (St. Louis, MO, USA), including metronidazole (MNZ), clarithromycin (CLR), amoxicillin (AMX), rifampicin (RIF), ciprofloxacin (CIP), levofloxacin (LEV), and tetracycline (TCN). The range of antibiotic concentrations was as follows: 0.25-256 mg/L for MNZ, 0.06 to 64 mg/L for CLR, 0.03 to 4 mg/L for AMX, 0.03 to 32 mg/L for RIF and LEV, 0.06 to 32 mg/L for CIP, and 0.06 to 16 mg/L for TCN. *H. pylori* inoculums were prepared from 72 h-old cultures that were suspended in sterile saline and adjusted to a density equal to No. 3 McFarland standard. The bacterial suspensions were inoculated directly onto Mueller-Hinton blood agar (Merck, Germany) plates supplemented with 10% defibrinated horse blood containing antibiotic dilutions, and were incubated under microaerophilic conditions as over-mentioned. After 72 hours of incubation, the minimal inhibition concentrations (MICs) were determined as the lowest concentration of antibiotic that completely inhibited the growth of the inoculums. The resistance breakpoints were used as described by the last guideline of European Committee on Antimicrobial Susceptibility Testing (EUCAST). Strains were considered to MNZ, >0.125 mg/L for AMX, and >1 mg/L for RIF, CIP, LEV, and TCN. Clarithromycin MICs were interpreted based on CLSI breakpoints (≤0.25 mg/L, susceptible; 0.5 mg/L; intermediate; ≥1.0 mg/L, resistant).^31^ A clinical isolate of *H. pylori* with previously identified MIC values was served as a quality control strain in all susceptibility tests.^32^

### Genomic DNA extraction

Subcultures of the single colonies were prepared, and confluent cultures from each colony were used for DNA extraction using QIAamp DNA extraction kit (QIAgen®, Hilden, Germany) following the manufacturer’s directions. The DNA samples were stored at −20 °C until used for gene amplification.

### Mutation analysis of the resistance genes

To detect specific mutations in the *frxA, rdxA, gyrA, gyrB* and 23S rRNA genes, a PCR-based sequencing approach was carried out in all *H. pylori* isolates including the susceptible and resistant strains. The *frxA* and *rdxA* genes were amplified as described by Han et al.^33^ Amplification of *gyrA* and *gyrB* genes were performed using primers as described elsewhere.^34^ Mutations within bacterial 23S rRNA peptidyl transferase gene were assessed as presented by Ho et al.^35^ The oligonucleotide primers are shown in Table 1. The PCR products were sequenced on both strands using an automated sequencer (Macrogen, Seoul, Korea). All complete and partial DNA sequences were edited by Chromas Lite version 2.5.1 (Technelysium Pty Ltd, Australia). Comparative sequence analysis between resistant and sensitive strains was carried out using BioEdit software version 7.2.5.^36^ The DNA and deduced amino acid sequences were aligned and coordinated to *H. pylori* 26695 (GenBank: CP003904.1) as a reference sequence.

**Table 1.**
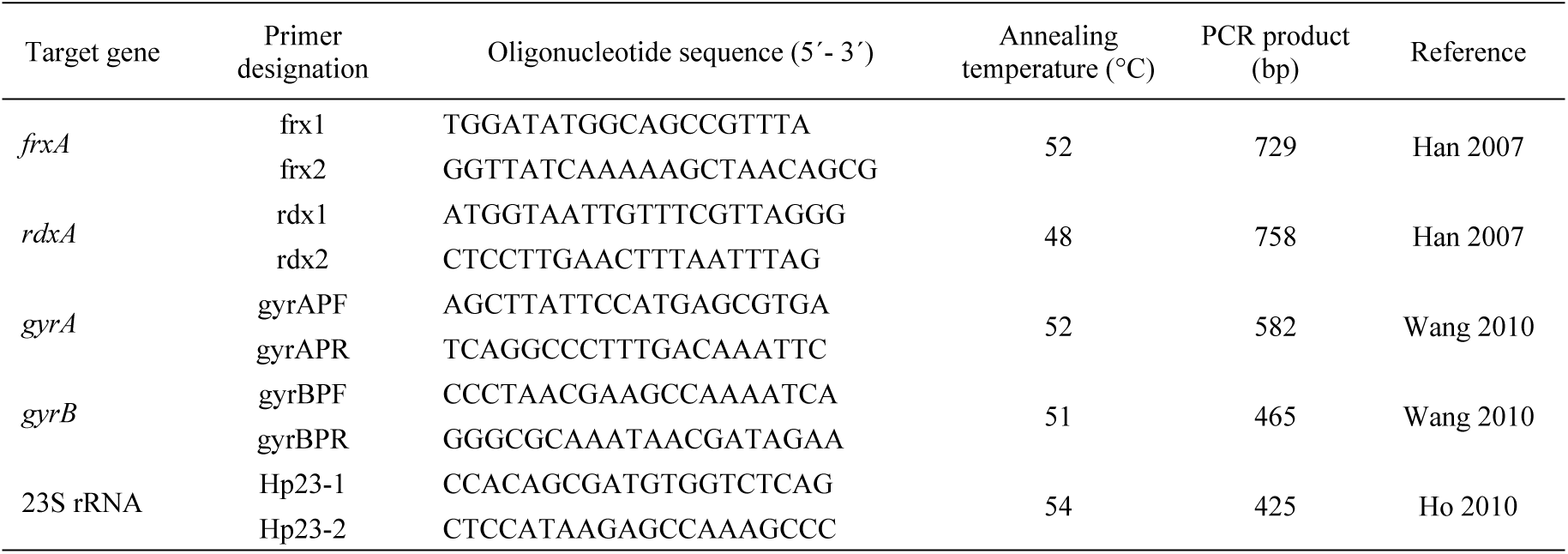
Oligonucleotide primer sequences used for amplification of genes involved in *H. pylori* antibiotic resistance.

### Detection of virulence markers

The presence of virulence factors including *cagA, cagL, vacA* alleles (s1/s2 and m1/m2), *babA2* and *sabA* genes were assessed based on our previously published work.^37^ The diversity of *cagA* C-terminal variable region and intactness of *cag*PAI was analysed as previously described.^38^ A PCR-sequencing assay was also used to analyse the functional (on/off) status of *oipA* gene.^29^ To investigate the presence of *dupA* gene, we used the previously designed primers by Jung *et al.*^39^ *H. pylori* J99 (CCUG 47164) and a no-template reaction were used as positive and negative controls in all amplifications, respectively.

Nucleotide sequence accession numbers

The sequences obtained from this study were submitted to NCBI under the following GenBank accession numbers: domain V 23S rRNA, MH040926-MH040949; *gyrA*, MH054262-MH054292; *gyrB*, MH054293-MH054319; *frxA*, MH054320-MH054346; *rdxA*, MH054347-MH054374.

### Statistical analysis

The SPSS Statistics for Windows (version 21.0, Armonk, NY: IBM Corp.) was used to perform all statistical analyses. The Chi-square and Fisher’s exact tests were used to determine the statistical significance of differences between categorical variables. A *P* value of less than 0.05 was considered as statistically significant.

## Results

### Characteristics of patients

Totally, 33 (42.3%) *H. pylori* isolates were cultured from antral biopsies of the patients included in the study. The *H. pylori* infected patients consisted of 14 (42.4%) men and 19 (57.6%) women, with an average age of 49.7 ± 9.7 years old (range 28-75 years). Endoscopic diagnosis showed that 15 patients had chronic gastritis (CG), 12 had peptic ulcer disease (PUD), and 6 had intestinal metaplasia (IM).

### Prevalence of antibiotic resistance

Overall, metronidazole resistance was the highest (27/33, 81.8%), and the lowest resistance rate was observed against tetracycline in 2/33 (6.1%) isolates. Resistance to clarithromycin, amoxicillin and rifampicin was observed in 12/33 (36.4%), 10/33 (30.3%), and 10/33 (30.3%) of isolates, respectively. Three (9.1%) isolates were found as intermediate to clarithromycin*. H. pylori* resistance to ciprofloxacin and levofloxacin was detected in 12/33 (36.4%), and 9/33 (27.3%) of isolates, respectively. Only one isolate was found to be susceptible to all antibiotics examined. The rate of resistance to metronidazole, clarithromycin, amoxicillin and levofloxacin was higher in patients with PUD and IM than with CG patients. Inversely, the rate of resistance to rifampicin was higher in CG patients than with PUD and IM. There were no important differences in the rate of resistance to ciprofloxacin and tetracycline between CG and with PUD and IM patients. All patients with IM were resistant to metronidazole. The MIC range, MIC50/MIC90, prevalence of resistance and distribution of MIC values for the *H. pylori* strains are shown in Tables 2 and 3.

**Table 2.** Distribution of the antibiotic resistance patterns, MIC range, MIC_50_ and MIC_90_ values for each antibiotic among *H. pylori* isolates used in this study.

| Antibiotic agents | MIC range | MIC<sub>50</sub> | MIC<sub>90</sub> | No. (%) of MIC (mg/L) |  |
| --- | --- | --- | --- | --- | --- |
|  |  |  |  | Susceptible | Resistance |
| MNZ | 0.25-128 | 32 | 128 | 6 (18.2) | 27 (81.8) |
| CLR<sup>a</sup> | 0.06-16 | 1 | 16 | 18 (54.5) | 12 (36.4) |
| AMX | 0.03-0.5 | 0.25 | 0.5 | 23 (69.7) | 10 (30.3) |
| CIP | 0.06-32 | 2 | 16 | 21 (63.6) | 12 (36.4) |
| LEV | 0.03-32 | 2 | 16 | 24 (72.7) | 9 (27.3) |
| RIF | 0.03-32 | 2 | 16 | 23 (69.7) | 10 (30.3) |
| TCN | 0.06-16 | 16 | >16 | 31 (93.9) | 2 (6.1) |

MNZ, metronidazole; CLR, clarithromycin; AMX, amoxicillin; CIP, ciprofloxacin; LEV, levofloxacin; RIF, rifampicin; TCN, tetracycline
<sup>a</sup>Three (9.1%) *H. pylori* isolates had intermediate susceptibility against clarithromycin based on CLSI breakpoints (MIC values equal to 0.5 mg/L)

**Table 3.** Distribution of MIC values for each antibiotic among *H. pylori* isolates used in this study.

| MIC (mg/L) | No. (%) |  |  |  |  |  |  |
| --- | --- | --- | --- | --- | --- | --- | --- |
|  | MNZ | CLR | AMX | CIP | LEV | RIF | TCN |
| 0.03 |  |  | 11 (33.3) |  | 3 (9.1) | 3 (9.1) |  |
| 0.06 |  | 10 (30.3) | 10 (30.3) | 1 (3) | 9 (27.3) | 5 (15.2) | 8 (24.2) |
| 0.12 |  | 6 (18.2) | 2 (6.1) | 7 (21.2) | 5 (15.2) | 6 (18.2) | 10 (30.3) |
| 0.25 | 3 (9.1) | 2 (6.1) | 5 (15.2) | 6 (18.2) | 1 (3) | 2 (6.1) | 5 (15.2) |
| 0.5 |  | 3 (9.1) | 5 (15.2) | 3 (9.1) | 5 (15.2) | 5 (15.2) | 5 (15.2) |
| 1 | 2 (6.1) | 3 (9.1) |  | 4 (12.1) | 1 (3) | 2 (6.1) | 3 (9.1) |
| 2 |  | 4 (12.1) |  | 4 (12.1) | 1 (3) | 3 (9.1) |  |
| 4 |  | 1 (3) |  | 1 (3) |  | 1 (3) |  |
| 8 | 2 (6.1) |  |  |  |  | 2 (6.1) |  |
| 16 | 7 (21.2) | 4 (12.1) |  | 6 (18.2) |  | 2 (6.1) |  |
| 32 | 6 (18.2) |  |  | 1 (3) |  | 2 (6.1) |  |
| 64 | 8 (24.2) |  |  |  |  |  |  |
| 128 | 5 (15.2) |  |  |  |  |  |  |
| 256 |  |  |  |  |  |  |  |

MNZ, metronidazole; CLR, clarithromycin; AMX, amoxicillin; CIP, ciprofloxacin; LEV, levofloxacin, RIF, rifampicin; TCN, tetracycline

### Multi-drug resistance

Single-drug resistance (SDR) was observed in 9 (27.3%) isolates, in which resistance to metronidazole was the most frequent SDR phenotype (5/9, 55.5%). Totally, 23/33 (69.7%) isolates showed multidrug resistance (MDR) phenotype, and 16 different MDR profiles were detected. No isolate was resistant to all tested antibiotics. The distribution of the SDR and MDR profiles within various clinical outcome groups is shown in Table 4. Resistance to MNZ + AMX and MNZ + RIF were equally the most common double-drug resistance profiles (2/6, 33.3%). Resistance to MNZ + CLR + CIP was found as the most frequent triple-drug resistance profile (3/11, 27.3%). All of the isolates from patients with IM and more than half of the PUD isolates showed MDR phenotype, mostly having triple-drug resistance profile.

**Table 4.** Distribution of the multidrug resistance profiles in relation to clinical outcomes among *H. pylori* isolates used in this study.

| Resistance profiles | Clinical outcome |  |  | Total<br>No. (%) |
| --- | --- | --- | --- | --- |
|  | CG<br>(n = 14) | PUD<br>(n = 12) | IM<br>(n = 6) |  |
| Single drugs |  |  |  |  |
| CLR | 0 | 1 | 0 | 1 (3.1) |
| AMX | 1 | 0 | 0 | 1 (3.1) |
| RIF | 1 | 0 | 0 | 1 (3.1) |
| CIP | 0 | 1 | 0 | 1 (3.1) |
| MNZ | 3 | 2 | 0 | 5 (15.6) |
| Double drugs |  |  |  |  |
| MNZ + AMX | 1 | 1 | 0 | 2 (6.2) |
| MNZ + RIF | 1 | 0 | 1 | 2 (6.2) |
| CIP + RIF | 1 | 0 | 0 | 1 (3.1) |
| MNZ + CIP | 0 | 1 | 0 | 1 (3.1) |
| Triple drugs |  |  |  |  |
| MNZ + CLR + LEV | 0 | 1 | 1 | 2 (6.2) |
| MNZ + AMX + RIF | 1 | 1 | 0 | 2 (6.2) |
| MNZ + CLR + CIP | 1 | 1 | 1 | 3 (9.4) |
| MNZ + CLR + AMX | 0 | 0 | 1 | 1 (3.1) |
| MNZ + AMX + CIP/LEV | 0 | 1 | 0 | 1 (3.1) |
| MNZ + RIF + CIP/LEV | 2 | 0 | 0 | 2 (6.2) |
| Quadruple drugs |  |  |  |  |
| MNZ + CLR + AMX + LEV | 0 | 1 | 0 | 1 (3.1) |
| MNZ + CLR + AMX + CIP/LEV | 0 | 0 | 1 | 1 (3.1) |
| MNZ + CLR + TCN + LEV | 1 | 0 | 0 | 1 (3.1) |
| MNZ + TCN + RIF + LEV | 0 | 0 | 1 | 1 (3.1) |
| MNZ + CLR + AMX + CIP | 0 | 1 | 0 | 1 (3.1) |
| MNZ + CLR + CIP + RIF | 1 | 0 | 0 | 1 (3.1) |

MNZ, metronidazole; CLR, clarithromycin; AMX, amoxicillin; CIP, ciprofloxacin; LEV, levofloxacin; RIF, rifampicin; TCN, tetracycline; CG, chronic gastritis; PUD, peptic ulcer disease; IM, intestinal metaplasia

### Genetic variations of *frxA* and *rdxA* genes

Totally, 27 of the *frxA* and 28 of the *rdxA* genes obtained from all isolates were sequenced and analysed as shown in Table 5. Fourteen (51.8%) isolates exhibiting resistance to metronidazole predominantly carried insertions and/or deletions resulting in translational frameshift mutations in the FrxA. One isolate was found to have stop codon at position Q5Stop resulting in premature termination codon (PTC), while missense mutations were found in 11/27 (40.7%) isolates. In addition, about one-third (10/28, 35.7%) of the isolates were found to have frameshift mutations in *rdxA* gene. Nonsense mutations resulting in PTC were identified in 3/28 (10.7%) isolates due to codon substitutions at position Q50Stop of RdxA. Missense mutations were distributed among 5 susceptible and 10 resistant isolates of the *rdxA* genes. One resistant strain had no mutation in both genes. The peptide sequence alignments for *frxA* and *rdxA* genes from metronidazole-susceptible and -resistant isolates in comparison with reference strain are presented in Supplementary Figs S1 and S2, respectively.

**Table 5.**
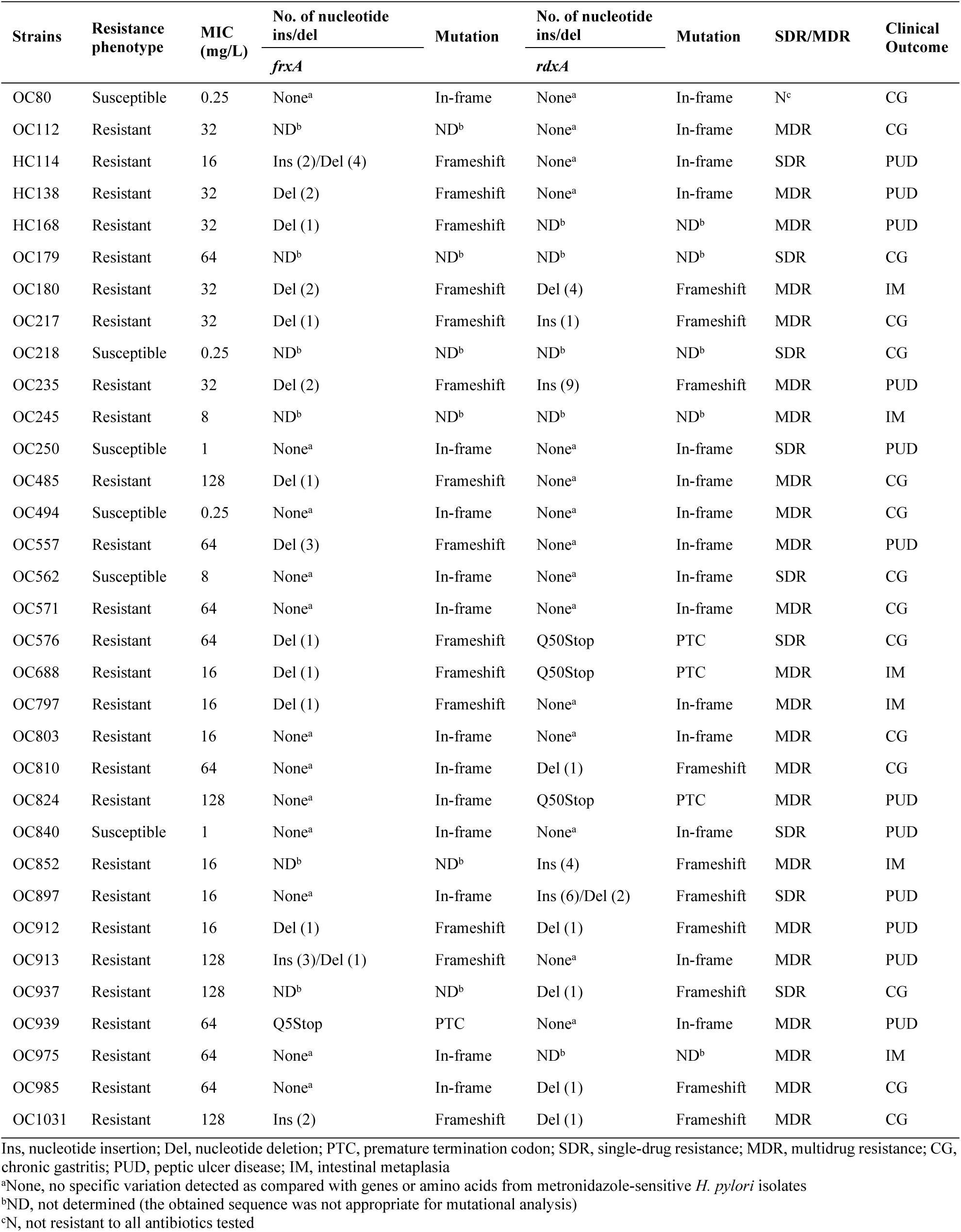
Number of nucleotide insertion and deletion in *frxA* and *rdxA* genes involved in metronidazole resistance among *H. pylori* isolates used in this study.

### Amino acid variations at QRDR region of *gyrA* and *gyrB* genes

As shown in Table 6 and Supplementary Figs S3 and S4, selective regions in the QRDR of *gyrA* and *gyrB* genes were sequenced among 31 and 27 *H. pylori* isolates, respectively. Totally, 16 different amino acid substitutions were detected in *gyrA* subunit among all isolates. Three different amino acid variants including S63P, R140K, and A183V were detected to be exclusively present in *gyrA* of the fluoroquinolone-resistant isolates, whereas six different substitutions of A97V, D143E, A207T, G208K, I212S, and E214K were found to be present in the susceptible isolates only. In addition, seven other mutations were observed at D86N, D86G, V150A, M191I, V199A, G208A, and G208E in both fluoroquinolone-susceptible and -resistant isolates. The most frequent substitutions in *gyrA* of the fluoroquinolone-resistant isolates were M191I (14/23, 60.9%), G208E (13/25, 52%), and V199A (5/9, 55.5%), respectively. The M191I-G208E substitution was found as the most common double mutations (8/12, 66.7%) within 5 resistant and 3 susceptible isolates, while the M191I-V199A-G208E was found as the most frequent triple substitutions (3/9, 33.3%) from 2 susceptible and one resistant isolates. The quadruple substitution was detected in *gyrA* of 3 resistant and 3 susceptible isolates.

**Table 6.** Mutations in *gyrA* and *gyrB* genes involved in fluoroquinolone resistance among *H. pylori* isolates used in this study.

| Strains | Resistance phenotype<br>CIP/LEV | MIC (mg/L) |  | Mutations |  | SDR/MDR | Clinical<br>Outcome |
| --- | --- | --- | --- | --- | --- | --- | --- |
|  |  | CIP | LEV | *gyrA* | *gyrB* |  |  |
| OC80 | Susceptible/Susceptible | 0.5 | 0.06 | M191I, V199A, G208E | D481E | N<sup>c</sup> | CG |
| OC112 | Susceptible/Susceptible | 0.25 | 0.25 | M191I, G208E | None<sup>a</sup> | MDR | CG |
| HC114 | Susceptible/Susceptible | 0.12 | 0.12 | M191I, G208E | None<sup>a</sup> | SDR | PUD |
| HC138 | Resistant/Susceptible | 16 | 0.5 | V150A, M191I, G208E | None<sup>a</sup> | MDR | PUD |
| HC168 | Susceptible/Susceptible | 0.25 | 0.06 | D143E, G208K, I212S, E214K | ND<sup>b</sup> | MDR | PUD |
| OC179 | Susceptible/Susceptible | 0.5 | 0.06 | M191I, V199A, G208A | ND<sup>b</sup> | SDR | CG |
| OC180 | Susceptible/Susceptible | 0.12 | 0.06 | V150A, M191I, V199A, G208E | None<sup>a</sup> | MDR | IM |
| OC217 | Susceptible/Susceptible | 0.12 | 0.06 | M191I, G208E | ND<sup>b</sup> | MDR | CG |
| OC218 | Susceptible/Susceptible | 0.12 | 0.06 | ND<sup>b</sup> | ND<sup>b</sup> | SDR | CG |
| OC235 | Susceptible/Susceptible | 0.25 | 0.06 | D86N, M191I, G208E | None<sup>a</sup> | MDR | PUD |
| OC245 | Resistant/Resistant | 16 | 16 | M191I, V199A, G208A | None<sup>a</sup> | MDR | IM |
| OC250<sup>d</sup> | Resistant/Susceptible | 16 | 0.5 | D86G, M191I | None<sup>a</sup> | SDR | PUD |
| OC485<sup>d</sup> | Resistant/Susceptible | 2 | 0.12 | D86N, A183V, M191I | None<sup>a</sup> | MDR | CG |
| OC494<sup>d</sup> | Resistant/Susceptible | 32 | 0.5 | None<sup>a</sup> | None<sup>a</sup> | MDR | CG |
| OC557 | Susceptible/Resistant | 0.12 | 16 | V199A, G208E | None<sup>a</sup> | MDR | PUD |
| OC562 | Susceptible/Susceptible | 1 | 0.12 | D86G, M191I, A207T, G208E | ND<sup>b</sup> | SDR | CG |
| OC571 | Resistant/Resistant | 16 | 16 | D86G, M191I, V199A, G208E | None<sup>a</sup> | MDR | CG |
| OC576 | Susceptible/Susceptible | 0.06 | 0.5 | V199A, G208E | D481E | SDR | CG |
| OC688 | Susceptible/Susceptible | 1 | 1 | M191I, V199A, G208E | None<sup>a</sup> | MDR | IM |
| OC797 | Resistant/Susceptible | 2 | 0.03 | M191I, V199A, G208E | D481E, R484K | MDR | IM |
| OC803 | Resistant/Susceptible | 2 | 0.03 | S63P, M191I, V199A, G208E | None<sup>a</sup> | MDR | CG |
| OC810 | Susceptible/Resistant | 1 | 32 | M191I, G208E | None<sup>a</sup> | MDR | CG |
| OC824 | Resistant/Resistant | 16 | 16 | M191I, G208E | D481E, R484K | MDR | PUD |
| OC840 | Susceptible/Susceptible | 1 | 0.06 | G208E | None<sup>a</sup> | SDR | PUD |
| OC852 | Susceptible/Resistant | 0.5 | 2 | M191I, G208E | D481E, R484K | MDR | IM |
| OC897 | Susceptible/Susceptible | 0.25 | 0.03 | A97V, G208E | None<sup>a</sup> | SDR | PUD |
| OC912 | Resistant/Susceptible | 2 | 0.12 | G208E | D481E, R484K | MDR | PUD |
| OC913 | Susceptible/Resistant | 0.25 | 16 | M191I, G208E | D481E, R484K | MDR | PUD |
| OC937 | Susceptible/Susceptible | 0.12 | 0.5 | G208E | None<sup>a</sup> | SDR | CG |
| OC939 | Resistant/Susceptible | 4 | 0.12 | M191I, G208E | None<sup>a</sup> | MDR | PUD |
| OC975 | Susceptible/Resistant | 0.25 | 16 | D86N, R140K, M191I, G208E | None<sup>a</sup> | MDR | IM |
| OC985 | Susceptible/Susceptible | 0.12 | 0.06 | ND<sup>b</sup> | None<sup>a</sup> | MDR | CG |
| OC1031 | Resistant/Resistant | 16 | 16 | D86N, M191I, G208E | ND<sup>b</sup> | MDR | CG |

548CIP, ciprofloxacin; LEV, levofloxacin; SDR, single-drug resistance; MDR, multidrug resistance; CG, chronic gastritis; PUD, peptic ulcer disease; IM, intestinal metaplasia
550<sup>a</sup>None, no specific variation detected as compared with genes or amino acids from fluoroquinolone-sensitive *H. pylori* isolates
551<sup>b</sup>ND, not determined (the obtained sequence was not appropriate for mutational analysis)
552<sup>c</sup>N, not resistant to all antibiotics tested
553<sup>d</sup>The *gyrA* quinolone-resistant determining regions of the strains OC250, OC485 and OC494 were partially translated due to low quality of obtained sequences

As for *gyrB* subunit, two different amino acid variants including D481E and R484K were detected among 7 isolates. The D481E substitution was found to be present in both fluoroquinolone-susceptible and -resistant isolates, whereas R484K was exclusively present in resistant isolates. As shown in Table 6 and Supplementary Fig S4, two fluoroquinolone-susceptible isolates had the single D481E mutations, while 5 resistant isolates presented the double D481E-R484K only. No mutation of *gyrB* was found in 11 fluoroquinolone-resistant and 9 susceptible isolates.

### Genetic variations of 23S rRNA gene

The domain V of 23S rRNA gene was sequenced in 24 *H. pylori* isolates. As shown in Supplementary Fig S5, this region was highly conserved with minimal nucleotide variations in comparison to *H. pylori* strain 26695 as the reference genome. Overall, four nucleotide transitions including A2143G and C2195T were identified in clarithromycin-resistant isolates. None of these mutations were observed among the susceptible isolates and no isolates were found to have double mutations of A2143G and C2195T. The distribution of MIC values according to the different mutations in all phenotypically resistant and susceptible isolates is presented in Table 7.

**Table 7.** Mutations in 23S rRNA gene involved in clarithromycin resistance among *H. pylori* isolates used in this study.

| Strains | Resistance phenotype | MIC (mg/L) | Mutations | SDR/MDR | Clinical Outcome |
| --- | --- | --- | --- | --- | --- |
| OC80 | Susceptible | 0.062 | None<sup>a</sup> | N<sup>c</sup> | CG |
| OC112 | Susceptible | 0.125 | ND<sup>b</sup> | MDR | CG |
| HC114 | Susceptible | 0.062 | None<sup>a</sup> | SDR | PUD |
| HC138 | Resistant | 16 | None<sup>a</sup> | MDR | PUD |
| HC168 | Susceptible | 0.062 | None<sup>a</sup> | MDR | PUD |
| OC179 | Susceptible | 0.062 | ND<sup>b</sup> | SDR | CG |
| OC180 | Resistant | 16 | A2143G | MDR | IM |
| OC217 | Susceptible | 0.062 | ND<sup>b</sup> | MDR | CG |
| OC218 | Susceptible | 0.25 | ND<sup>b</sup> | SDR | CG |
| OC235 | Intermediate | 0.5 | None<sup>a</sup> | MDR | PUD |
| OC245 | Resistant | 16 | ND<sup>b</sup> | MDR | IM |
| OC250 | Susceptible | 0.25 | None<sup>a</sup> | SDR | PUD |
| OC485 | Resistant | 2 | None<sup>a</sup> | MDR | CG |
| OC494 | Susceptible | 0.062 | None<sup>a</sup> | MDR | CG |
| OC557 | Resistant | 2 | C2195T | MDR | PUD |
| OC562 | Intermediate | 0.5 | ND<sup>b</sup> | SDR | CG |
| OC571 | Susceptible | 0.25 | None<sup>a</sup> | MDR | CG |
| OC576 | Susceptible | 0.125 | ND<sup>b</sup> | SDR | CG |
| OC688 | Susceptible | 0.125 | None<sup>a</sup> | MDR | IM |
| OC797 | Resistant | 16 | None<sup>a</sup> | MDR | IM |
| OC803 | Resistant | 1 | ND<sup>b</sup> | MDR | CG |
| OC810 | Resistant | 2 | C2195T | MDR | CG |
| OC824 | Susceptible | 0.062 | None<sup>a</sup> | MDR | PUD |
| OC840 | Resistant | 1 | A2143G | SDR | PUD |
| OC852 | Resistant | 2 | ND<sup>b</sup> | MDR | IM |
| OC897 | Susceptible | 0.062 | None<sup>a</sup> | SDR | PUD |
| OC912 | Intermediate | 0.5 | None<sup>a</sup> | MDR | PUD |
| OC913 | Susceptible | 0.125 | None<sup>a</sup> | MDR | PUD |
| OC937 | Susceptible | 0.125 | None<sup>a</sup> | SDR | CG |
| OC939 | Resistant | 4 | None<sup>a</sup> | MDR | PUD |
| OC975 | Susceptible | 0.125 | None<sup>a</sup> | MDR | IM |
| OC985 | Susceptible | 0.125 | None<sup>a</sup> | MDR | CG |
| OC1031 | Susceptible | 0.062 | None<sup>a</sup> | MDR | CG |

SDR, single-drug resistance; MDR, multidrug resistance; CG, chronic gastritis; PUD, peptic ulcer disease; IM, intestinal metaplasia
<sup>a</sup>None, no specific variation detected as compared with genes from clarithromycin-sensitive *H. pylori* isolates
<sup>b</sup>ND, not determined (the obtained sequence was not appropriate for mutational analysis)
<sup>c</sup>N, not resistant to all antibiotics tested

### Association between virulence genotypes and resistance patterns

The frequency and distribution of strains grouped by virulence genotypes according to each susceptibility pattern is shown in Table 8. High frequencies of *cagL*-positive and *cagA*-positive genotypes were found frequently in susceptible isolates for all antibiotics tested, with the exception of metronidazole. The *cagA* ABC motif was also found frequently in susceptible isolates, with the exception of metronidazole and clarithromycin. *H. pylori* isolates with *vacA* s1m2 were found more frequently in susceptible isolates. Strains with *oipA* “on” status and *babA, sabA* and *dupA* positivity were also frequently found in susceptible isolates. There were only two isolates that showed resistance against tetracycline and both these strains also were *sabA*-positive. All ciprofloxacin-resistant isolates also were found to be *dupA*-positive. There was no association between these virulence factors and antibiotic resistance patterns (*P* > 0.05). Furthermore, *H. pylori* strains harboring intact or partial *cag*PAI were variably distributed between susceptible and resistant isolates. Interestingly, significant associations were observed between intact *cag*PAI and resistance to rifampicin (*P* = 0.027), and between susceptibility to amoxicillin and *cag*PAI intactness (*P* = 0.016).

**Table 8.**
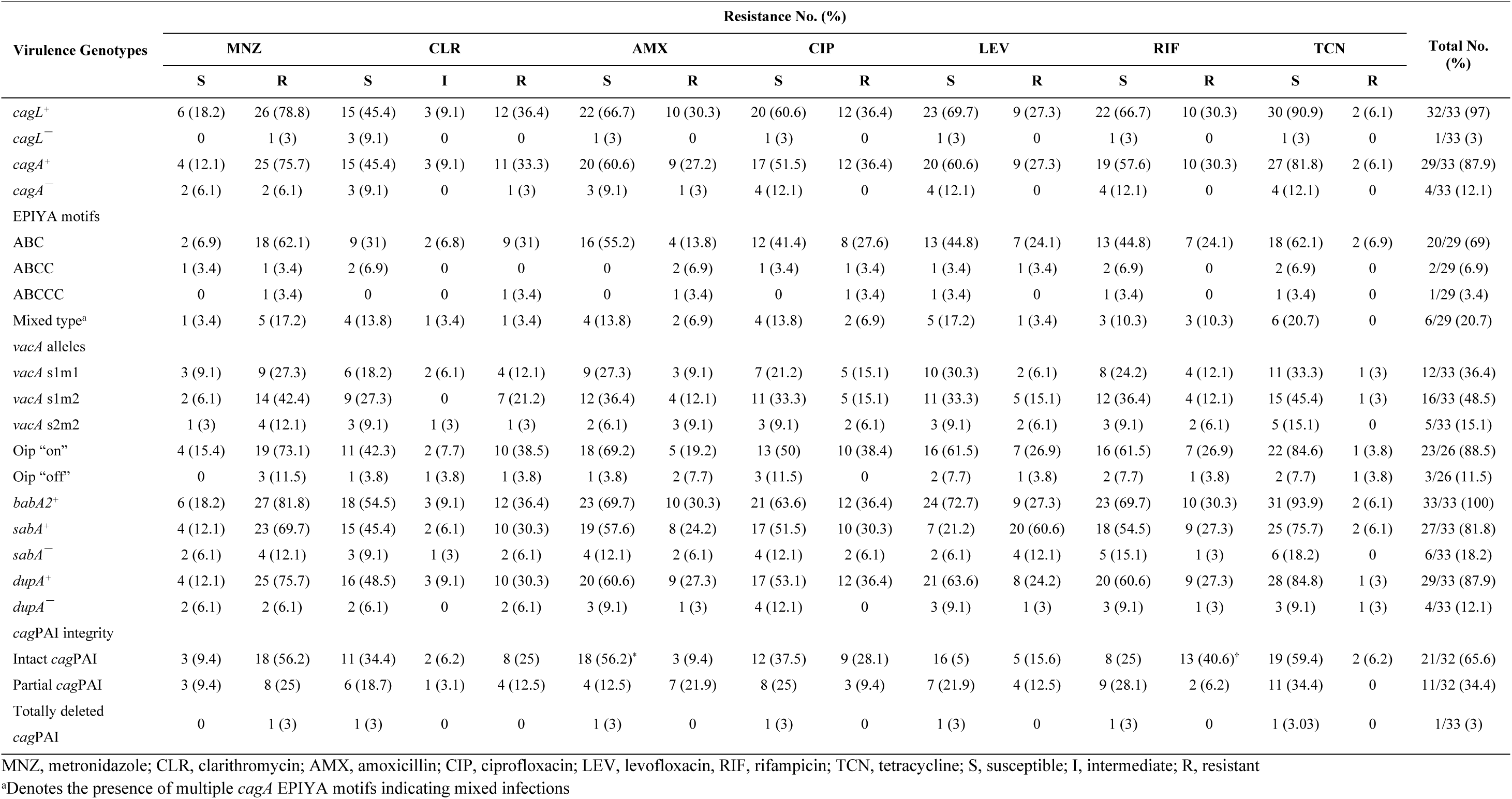
Frequency and distribution of virulence genotypes in relation to antibiotic resistance patterns among *H. pylori* isolates used in this study.

## Discussion

Eradication of *H. pylori* infection has been reported to significantly improve the clinical outcome of infected patients in high-risk areas.^3,40^ However, the eradication rate of *H. pylori* has been decreasing progressively, mainly due to increased resistance to antimicrobial agents, especially in developing countries.^5,6,11,17,41-43^ Thus, in order to choose the appropriate antibiotics in different *H. pylori* treatment regimens, we need to have recently updated susceptibility data in the local setting. In Iran, nearly 40-90% of the adult population is infected with *H. pylori*, which seems to be acquired early in childhood.^32,37,44^ There have been few reports on the antibiotic resistance of *H. pylori* in Iran by performing agar dilution method as the reference method for this bacterium.^32,45-47^ However, none of the previous studies investigated the functional and molecular mechanisms that contribute to resistance of *H. pylori* strains from Iranian patients. Therefore, we carried out this work to determine the molecular characteristics of genes involved in antibiotic resistance, and evaluate the association between resistance patterns and a wide panel of virulence genotypes. The prevalence of metronidazole resistance was reported to be high among Iranian *H. pylori* strains and ranged from 40.5% to 78.6%.^45,47,48^ The results of this study showed an increased rate of metronidazole resistance (81.8%) as compared to previous reports from Iran.^32,45,47^ A very high prevalence of metronidazole resistance has also been reported from other developing countries in Asia including Bangladesh (77.5%), China (95.4%), India (83.8%), Kuwait (70%), Pakistan (89%), and Vietnam (69.9%).^43,49-53^ The extremely high rate of metronidazole resistance observed in this study might be attributed to widespread and unauthorized consumption of antimicrobial drugs in Iran. In addition, massive use of metronidazole in the treatment of various infections such as anaerobic bacterial and parasitic infections, and for diarrheal, dental, periodontal, and gynecologic diseases could explain the significantly high rate of metronidazole resistance in in many developing countries.^5,32,41,43^ Therefore, in agreement with other previous studies *H. pylori* treatment regimens containing metronidazole are not useful and should not be chosen as first-line eradication therapy in Iran.^5,41-43,54^

Previous studies demonstrated that various point mutations in *frxA* and *rdxA* genes were linked to metronidazole resistance in *H. pylori*.^23,51,55^ As expected, different types of mutations including insertions, deletions, missense, nonsense and frameshift mutations were detected among the studied strains. Our results showed that most of the metronidazole-resistant isolates presented frameshift mutations in these genes. Moreover, we found point mutations introducing stop codon at positions Q5Stop and Q50Stop in *frxA* and *rdxA* genes, respectively. Many other nonsense mutations that lead to PTC have also been reported in *rdxA* and rarely in *frxA* genes.^5,17,22,33,41,56,57^ However, in this study one of the metronidazole-resistant isolate did not contain any alterations in both *frxA* and *rdxA* genes. As previously suggested, metronidazole resistance in this small subset of isolates may be due to the presence of additional resistance mechanisms and mutations in other redox enzymes.^33,41,58^

Fluoroquinolones were proven to have bacteriostatic activities by trapping DNA gyrase and topoisomerase IV. These drugs considered as salvage treatment for *H. pylori* eradication in second-or third-line therapies after the failure of clarithromycin-based treatment regimens.^12,24,59^ However, it has been reported that fluoroquinolone resistance is rapidly expanding around the world.^6,7,53^ In a previous study from Iran, the rate of resistance to ciprofloxacin and levofloxacin was reported about 27% and 24.3%, respectively.^32^ In this study, we found a significant increase of fluoroquinolone resistance, which is of great concern. Nevertheless, studies from Taiwanese and Malaysian populations revealed that gemifloxacin is superior to levofloxacin in antimicrobial activity and may have better drug efficacy than levofloxacin in *H. pylori* eradication.^41,60^

Point mutations in the QRDR of *gyrA* and *gyrB* sequences, greatly reduce the antimicrobial activity of fluoroquinolones. To date, several mutations have been identified in *gyrA* subunit of *H. pylori* strains from different geographical regions.^5,8,34,41,60-64^ None of the most common mutations in *gyrA* hot spot positions N87K, D91N, and D91G were detected in our isolates. However, 10 novel substitutions including S63P, D143E, A183V, A207T, G208K, G208A, G208E, I212S, E214K, and M191I were identified in the *gyrA* of either/both fluoroquinolone-resistant or/and -susceptible strains in this study. Among them mutations M191I, G208E, and V199A were predominantly found in fluoroquinolone-resistant isolates. Moreover, *gyrB* mutations may rarely occur and have little impact on primary fluoroquinolone resistance.^5,8,34,41,61,64^ In this study, only two amino acid changes D481E and R484K were identified in *gyrB*, in which R484K was exclusively present in resistant strains.

Among macrolides, clarithromycin is recognized as a major antibiotic for *H. pylori* eradication therapy because of its impact on treatment outcomes.^20,65^ The rate of clarithromycin resistance is topically much lower than that to metronidazole. However, the rate of primary clarithromycin resistance is undoubtedly on the rise and varies between different geographical regions.^7,8,43,53,66^ Unfortunately, the level of clarithromycin resistance in this study increased in comparison to a previous study^32^ from 26 to 36.4%, which is of great concern.

It has been claimed that three most frequently reported mutations, including A2143G, A2142G and A2142C, are responsible for more than 90% cases of primary resistance to clarithromycin.^67,68^ However, in a recent study by De Francesco *et al.* this concordance was reduced to only 54.8%, with the A2142C mutation not being detected at all.^20^ Moreover, some other mutations have been found to be associated with clarithromycin resistance, although their precise role is not yet clear.^69^ In this study, only four mutations including A2143G and C2195T were found in our isolates. We failed to identify additional mutations such as T2183C and A2223G, which are frequently reported to be the cause of clarithromycin resistance in Eastern countries, rather than in Western countries.^70^ Additionally, no point mutation was identified in the sequence of 23S rRNA gene in four clarithromycin-resistant strains. For those isolates, we can speculate that other resistance mechanisms, such as the presence of an efflux pump, may be implicated in development of resistance to clarithromycin.^71^

It is estimated that the overall resistance rates to amoxicillin and tetracycline are 23.61% and 7.38% in Asian countries, respectively.^48^ Similarly, we observed high rate of resistance to these drugs among the studied isolates (30.3% to amoxicillin and 6.1% to tetracycline), which is a matter of great concern in *H. pylori* eradication in Iran. However, the level of resistance to these antibiotics reported to be very low or even absent in most western countries versus African countries.^69,72^ Regarding rifampicin, we also observed a rising rate of resistance from 14.4% to 30.3% in comparison to a previous report.^32^ Recently, Regnath *et al.* reported considerable increase in resistance to rifampicin from 3.9% to 18.8% between 2002 and 2015 among pediatric patients from southwest Germany.^73^

Unfortunately, emergence of MDR *H. pylori* strains has become a serious challenge all over the world. In a previous study, the resistance rate to at least two antimicrobial agents was reported in 43% of the *H. pylori* isolates from Iran. Surprisingly, our finding showed that 69.7% of the isolates were resistant to at least two antibiotics. The high prevalence of MDR phenotype may be attributed to the exhaustive use of antibiotics across the country. Information about the prevalence of quadruple-drug resistance is limited, and a few reports from India (2.5%), Bulgaria (0.7%), Vietnam (1.9%) and Indonesia (2.6%) are available yet.^5,43,74,75^ However, 18.7% of the isolates in this study showed quadruple-drug resistance, which was lower than the previous study (37.9%).^32^ Moreover, resistance to tetracycline was only observed in the isolates with quadruple-drug resistance. This finding could be explained to the presence of multidrug efflux pumps in these strains.^69^

There have been several reports on the relationship between *H. pylori* virulence markers and antibiotic resistance. Accordingly, patients infected with *cagA*-positive strains that also carry more virulent *vacA* alleles significantly have high cure rates and eradication success than less virulent strains.^26,76-78^ It has been hypothesized that colonization of gastric mucosa by more virulent *H. pylori* genotypes may induce a higher degree of inflammation and increase blood flow, which in turn can favor better diffusion of the antibiotics.^27,78^ Alternatively, another possible explanation may be related to the fact that *cagA*-positive strains proliferate faster than *cagA*-negative ones and would therefore be more susceptible to antibiotics.^25,76^ Furthermore, Taneike *et al.* observed that *cagA*-negative strains may tend to acquire spontaneous drug resistance under selective pressure of antimicrobials.^25^ However, it still remains somewhat controversial because recent reports indicated that these virulent genotypes variously distributed between susceptible and resistant strains.^5,28,79-81^ CagA protein with a greater number of EPIYA-C repeats is considered to be pathophysiologically more virulent and carcinogenic.^82^ Thus, according to the over-mentioned hypothesis, we expected the presence of more virulent types of CagA EPIYA motifs in susceptible isolates than resistant ones. However, as the number of EPIYA types having two or more EPIYA-C repeats was very low, we could not come to such conclusion. Similar to other studies performed in Italy (37.2%)^83^, North Wales (53%)^84^ and Germany (37.4%)^77^, *vacA* s1m2 (48.5%) genotype was the most prevalent *vacA* mosaicisms in our strains. Although *H. pylori* strains with *vacA* s1m2 were detected more frequently in metronidazole-resistant isolates, no significant associations was found. In contrast, *vacA* s1m2 genotype was found more frequently in susceptible isolates for other antibiotics examined. Likewise, isolates with *oipA* “on” status and harboring *babA, sabA* and *dupA* genotypes were frequently found in metronidazole-resistant isolates. On the other hand, all of these genotypes were frequently found in susceptible isolates for other antibiotics, with the exception of *sabA* in levofloxacin-resistant ones. Moreover, two isolates that showed resistance against tetracycline were *sabA*-positive, and all ciprofloxacin-resistant isolates were found to be *dupA*-positive. However, we found no association between these virulence factors and antibiotics resistance (*P* > 0.05). *H. pylori* strains that carry an intact and functional *cag*PAI are more virulent and frequently associated with severe clinical outcomes than those carrying partial or no *cag*PAI.^30,38^ As far as we know, this is the first study that relates the *cag*PAI integrity with antibiotic resistance. Our results showed that *H. pylori* isolates harboring intact or partial *cag*PAI were variably distributed between susceptible and resistant isolates. However, we found significant associations between intact *cag*PAI and resistance to rifampicin (*P* = 0.027), and contrastingly between susceptibility to amoxicillin and *cag*PAI intactness (*P* = 0.016). These results are contradictory and did not strongly support the idea that susceptibility to antibiotics is higher in infections caused by more virulent genotypes. Nevertheless, it is likely that infected patients with resistant and hypervirulent strains are at increased risk of progression to more severe clinical outcomes due to failure in *H. pylori* eradication.

In conclusion, this study demonstrated that the prevalence of *H. pylori* antibiotic resistance is worrisome in our country with rising trend over the time. The findings from this study also highlight the relevance of different types of mutations in genes responsible for antibiotic resistance in *H. pylori* strains. We also provide evidence for the importance of simultaneous screening of the virulence and resistance genotypes in *H. pylori* strains for guiding clinicians to choose an appropriate combination of drugs. Taken together, because of alarming increase in the rate of *H. pylori* antibiotic resistance in our local population, it is reasonable to constantly monitor the antimicrobial susceptibility patterns, and develop effective treatment and preventive strategies at national level.

## Acknowledgements

This study was supported by a grant (no. RIGLD 878) from Research Institute for Gastroenterology and Liver Diseases, Shahid Behehsti University of Medical Sciences, Tehran, Iran.

## Author Contributions

N. Farzi collected the *H. pylori* strains, performed the susceptibility testing and molecular assays. A. Yadegar worked on concept and design of the study, data analysis and interpretation, and writing of manuscript. A. Sadeghi, H. Asadzadeh Aghdaei, and M. R. Zali critically revised the paper. All authors approved the final version of the manuscript and the authorship list.

## Supplementary Information

Supplementary information associated with this article can be found, in the online version.

## References

1. Backert S, Neddermann M, Maubach G et al. Pathogenesis of *Helicobacter pylori* infection. Helicobacter 2016;21 Suppl 1:19–25.

2. Rowland M, Daly L, Vaughan M et al. Age-specific incidence of *Helicobacter pylori*. Gastroenterology 2006;130:65-72; quiz 211.

3. Malfertheiner P, Megraud F, O’Morain CA et al. Management of *Helicobacter pylori* infection-the Maastricht V/Florence Consensus Report. Gut 2017;66:6–30.

4. Wu CY, Kuo KN, Wu MS et al. Early *Helicobacter pylori* eradication decreases risk of gastric cancer in patients with peptic ulcer disease. Gastroenterology 2009;137:1641–48.

5. Miftahussurur M, Syam AF, Nusi IA et al. Surveillance of *Helicobacter pylori* antibiotic susceptibility in Indonesia: different resistance types among regions and with novel genetic mutations. PloS One 2016;11:e0166199.

6. Ngoyi O, Nina E, Atipo Ibara BI et al. Molecular detection of *Helicobacter pylori* and its antimicrobial resistance in Brazzaville, Congo. Helicobacter 2015;20:316–20.

7. Lee JW, Kim N, Kim JM et al. Prevalence of primary and secondary antimicrobial resistance of *Helicobacter pylori* in Korea from 2003 through 2012. Helicobacter 2013;18:206–14.

8. Zerbetto De Palma G, Mendiondo N, Wonaga A et al. Occurrence of mutations in the antimicrobial target genes related to levofloxacin, clarithromycin, and amoxicillin resistance in *Helicobacter pylori* isolates from Buenos Aires city. Microb Drug Resist 2017;23:351–8.

9. Mahachai V, Vilaichone RK, Pittayanon R et al. Helicobacter pylori management in ASEAN: The Bangkok consensus report. J Gastroenterol Hepatol 2018;33:37–56.

10. Mégraud F. The challenge of Helicobacter pylori resistance to antibiotics: the comeback of bismuth-based quadruple therapy. Therap Adv Gastroenterol 2012;5:103–9.

11. Kuo CH, Hsu PI, Kuo FC et al. Comparison of 10 day bismuth quadruple therapy with high-dose metronidazole or levofloxacin for second-line *Helicobacter pylori* therapy: a randomized controlled trial. J Antimicrob Chemother 2013;68:222–8.

12. Gisbert JP, Romano M, Gravina AG et al. Helicobacter pylori second-line rescue therapy with levofloxacin- and bismuth-containing quadruple therapy, after failure of standard triple or non-bismuth quadruple treatments. Aliment Pharmacol Ther 2015;41:768–75.

13. Cianci R, Montalto M, Pandolfi F et al. Third-line rescue therapy for *Helicobacter pylori* infection. World J Gastroenterol 2006;12:2313–19.

14. Delchier JC, Malfertheiner P, Thieroff-Ekerdt R. Use of a combination formulation of bismuth, metronidazole and tetracycline with omeprazole as a rescue therapy for eradication of Helicobacter pylori. Aliment Pharmacol Ther 2014;40:171–7.

15. Chen Q, Zhang W, Fu Q, et al. Rescue therapy for *Helicobacter pylori* eradication: a randomized non-inferiority trial of amoxicillin or tetracycline in bismuth quadruple therapy. Am J Gastroenterol 2016;111:1736–42.

16. Miftahussurur M, Shrestha PK, Subsomwong P et al. Emerging *Helicobacter pylori* levofloxacin resistance and novel genetic mutation in Nepal. BMC Microbiol 2016;16:256.

17. Alfizah H, Norazah A, Hamizah R et al. Resistotype of *Helicobacter pylori* isolates: the impact on eradication outcome. J Med Microbiol 2014;63:703–9.

18. Secka O, Berg DE, Antonio M et al. Antimicrobial susceptibility and resistance patterns among *Helicobacter pylori* strains from the Gambia, West Africa. Antimicrob Agents Chemother 2013;57:1231–7.

19. Nishizawa T, Suzuki H. Mechanisms of Helicobacter pylori antibiotic resistance and molecular testing. Front Mol Biosci 2014;1:19.

20. De Francesco V, Zullo A, Giorgio F et al. Change of point mutations in *Helicobacter pylori* rRNA associated with clarithromycin resistance in Italy. J Med Microbiol 2014;63:453–7.

21. Rimbara E, Noguchi N, Kawai T et al. Novel mutation in 23S rRNA that confers low-level resistance to clarithromycin in *Helicobacter pylori*. Antimicrob Agents Chemother 2008;52:3465–6.

22. Matteo MJ, Perez CV, Domingo MR et al. DNA sequence analysis of *rdxA* and *frxA* from paired metronidazole-sensitive and -resistant *Helicobacter pylori* isolates obtained from patients with heteroresistance. Int J Antimicrob Agents 2006;27:152–8.

23. Masaoka T, Suzuki H, Kurabayashi K et al. Could frameshift mutations in the *frxA* and *rdxA* genes of *Helicobacter pylori* be a marker for metronidazole resistance? Aliment Pharmacol Ther symp ser 2006;2:81–7.

24. Rimbara E, Noguchi N, Kawai T, Sasatsu M. Fluoroquinolone resistance in Helicobacter pylori: role of mutations at position 87 and 91 of GyrA on the level of resistance and identification of a resistance conferring mutation in *GyrB*. Helicobacter 2012;17:36–42.

25. Taneike I, Nami A, O’Connor A et al. Analysis of drug resistance and virulence-factor genotype of Irish *Helicobacter pylori* strains: is there any relationship between resistance to metronidazole and *cagA* status? Aliment Pharmacol Ther 2009;30:784–90.

26. Suzuki T, Matsuo K, Sawaki A et al. Systematic review and meta-analysis: importance of CagA status for successful eradication of *Helicobacter pylori* infection. Aliment Pharmacol Ther 2006;24:273–80.

27. Agudo S, Perez-Perez G, Alarcon T et al. High prevalence of clarithromycin-resistant *Helicobacter pylori* strains and risk factors associated with resistance in Madrid, Spain. J Clin Microbiol 2010;48:3703–7.

28. Fasciana T, Cala C, Bonura C et al. Resistance to clarithromycin and genotypes in *Helicobacter pylori* strains isolated in Sicily. J Med Microbiol 2015;64:1408–14.

29. Farzi N, Yadegar A, Aghdaei HA et al. Genetic diversity and functional analysis of *oipA* gene in association with other virulence factors among *Helicobacter pylori* isolates from Iranian patients with different gastric diseases. Infect Genet Evol 2018;60:26–34.

30. Ahmadzadeh A, Ghalehnoei H, Farzi N, et al. Association of *Cag*PAI integrity with severeness of *Helicobacter pylori* infection in patients with gastritis. Pathol Biol 2015;63:252–7.

31. Clinical and Laboratory Standards Institute. Methods for antimicrobial dilution and disk susceptibility testing of infrequently isolated or fastidious bacteria-3rd ed. Document M45. CLSI, Wayne, PA, USA, 2015.

32. Shokrzadeh L, Alebouyeh M, Mirzaei T et al. Prevalence of multiple drug-resistant *Helicobacter pylori* strains among patients with different gastric disorders in Iran. Microb Drug Resist 2015;21:105–10.

33. Han F, Liu S, Ho B et al. Alterations in *rdxA* and *frxA* genes and their upstream regions in metronidazole-resistant *Helicobacter pylori* isolates. Res Microbiol 2007;158:38–44.

34. Wang L-H, Cheng H, Hu F-L, Li J. Distribution of gyrA mutations in fluoroquinolone-resistant Helicobacter pylori strains. World J Gastroenterol 2010;16:2272–7.

35. Ho SL, Tan EL, Sam CK et al. Clarithromycin resistance and point mutations in the 23S rRNA gene in *Helicobacter pylori* isolates from Malaysia. J Dig Dis 2010;11:101–5.

36. Hall TA. BioEdit: a user-friendly biological sequence alignment editor and analysis program for Windows 95/98/NT. Nucl Acids Symp Ser 1999;41;95–8.

37. Yadegar A, Mobarez AM, Alebouyeh M et al. Clinical relevance of *cagL* gene and virulence genotypes with disease outcomes in a *Helicobacter pylori* infected population from Iran. World J Microbiol Biotechnol 2014;30:2481–90.

38. Yadegar A, Alebouyeh M, Zali MR. Analysis of the intactness of *Helicobacter pylori cag* pathogenicity island in Iranian strains by a new PCR-based strategy and its relationship with virulence genotypes and EPIYA motifs. Infect Genet Evol 2015;35:19–26.

39. Jung SW, Sugimoto M, Shiota S et al. The intact *dupA* cluster is a more reliable *Helicobacter pylori* virulence marker than *dupA* alone. Infect Immun 2012;80:381–7.

40. Lee YC, Chiang TH, Chou CK et al. Association between *Helicobacter pylori* eradication and gastric cancer incidence: a systematic review and meta-analysis. Gastroenterology 2016;150:1113–24.

41. Teh X, Khosravi Y, Lee WC et al. Functional and molecular surveillance of *Helicobacter pylori* antibiotic resistance in Kuala Lumpur. PloS One 2014;9:e101481.

42. Goh KL, Navaratnam P. High Helicobacter pylori resistance to metronidazole but zero or low resistance to clarithromycin, levofloxacin, and other antibiotics in Malaysia. Helicobacter 2011;16:241–5.

43. Binh TT, Shiota S, Nguyen LT et al. The incidence of primary antibiotic resistance of *Helicobacter pylori* in Vietnam. J Clin Gastroenterol 2013;47:233–8.

44. Hosseini E, Poursina F, de Wiele TV et al. Helicobacter pylori in Iran: a systematic review on the association of genotypes and gastroduodenal diseases. J Res Med Sci 2012;17:280–92.

45. Talebi Bezmin Abadi A, Ghasemzadeh A, Taghvaei T et al. Primary resistance of *Helicobacter pylori* to levofloxacin and moxifloxacine in Iran. Intern Emerg Med 2012;7:447–52.

46. Kohanteb J, Bazargani A, Saberi-Firoozi M et al. Antimicrobial susceptibility testing of *Helicobacter pylori* to selected agents by agar dilution method in Shiraz-Iran. Indian J Med Microbiol 2007;25:374–7.

47. Shokrzadeh L, Jafari F, Dabiri H et al. Antibiotic susceptibility profile of *Helicobacter pylori* isolated from the dyspepsia patients in Tehran, Iran. Saudi J Gastroenterol 2011;17:261–4.

48. Ghotaslou R, Leylabadlo HE, Asl YM. Prevalence of antibiotic resistance in *Helicobacter pylori*: A recent literature review. World J Methodol 2015;5:164–74.

49. Nahar S, Mukhopadhyay AK, Khan R et al. Antimicrobial susceptibility of *Helicobacter pylori* strains isolated in Bangladesh. J Clin Microbiol 2004;42:4856–8.

50. Pandya HB, Agravat HH, Patel JS et al. Emerging antimicrobial resistance pattern of *Helicobacter pylori* in central Gujarat. Indian J Med Microbiol 2014;32:408–13.

51. John Albert M, Al-Mekhaizeem K, Neil L et al. High prevalence and level of resistance to metronidazole, but lack of resistance to other antimicrobials in *Helicobacter pylori*, isolated from a multiracial population in Kuwait. Aliment Pharmacol Ther 2006;24:1359–66.

52. Khan A, Farooqui A, Manzoor H, Akhtar SS, Quraishy MS, Kazmi SU. Antibiotic resistance and *cagA* gene correlation: a looming crisis of *Helicobacter pylori*. World J Gastroenterol 2012;18:2245–52.

53. Su P, Li Y, Li H et al. Antibiotic resistance of *Helicobacter pylori* isolated in the southeast coastal region of China. Helicobacter 2013;18:274-9.

54. Ahmad N, Zakaria WR, Mohamed R. Analysis of antibiotic susceptibility patterns of *Helicobacter pylori* isolates from Malaysia. Helicobacter 2011;16:47–51.

55. Gerrits MM, Van der Wouden E-J et al. Role of the *rdxA* and *frxA* genes in oxygen-dependent metronidazole resistance of *Helicobacter pylori*. J Med Microbiol 2004;53:1123–8.

56. Marais A, Bilardi C, Cantet F et al. Characterization of the genes *rdxA* and *frxA* involved in metronidazole resistance in *Helicobacter pylori*. Res Microbiol 2003;154:137–44.

57. Yang YJ, Wu JJ, Sheu BS et al. The *rdxA* gene plays a more major role than *frxA* gene mutation in high-level metronidazole resistance of *Helicobacter pylori* in Taiwan. Helicobacter 2004;9:400–7.

58. 58 Chisholm SA, Owen RJ. Mutations in *Helicobacter pylori rdxA* gene sequences may not contribute to metronidazole resistance. J Antimicrob Chemother 2003;51:995–9.

59. Goh KL, Manikam J, Qua CS. High-dose rabeprazole-amoxicillin dual therapy and rabeprazole triple therapy with amoxicillin and levofloxacin for 2 weeks as first and second line rescue therapies for *Helicobacter pylori* treatment failures. Aliment Pharmacol Ther 2012;35:1097–102.

60. Chang WL, Kao CY, Wu CT et al. Gemifloxacin can partially overcome quinolone resistance of *H. pylori* with *gyrA* mutation in Taiwan. Helicobacter 2012;17:210–5.

61. Trespalacios-Rangel AA, Otero W et al. Surveillance of levofloxacin resistance in *Helicobacter pylori* isolates in Bogota-Colombia (2009-2014). PloS One 2016;11:e0160007.

62. Cattoir V, Nectoux J, Lascols C, et al. Update on fluoroquinolone resistance in *Helicobacter pylori*: new mutations leading to resistance and first description of a *gyrA* polymorphism associated with hypersusceptibility. Int J Antimicrob Agents 2007;29:389–96.

63. Garcia M, Raymond J, Garnier M et al. Distribution of spontaneous *gyrA* mutations in 97 fluoroquinolone-resistant *Helicobacter pylori* isolates collected in France. Antimicrob Agents Chemother 2012;56:550–1.

64. Miyachi H, Miki I, Aoyama N et al. Primary levofloxacin resistance and *gyrA/B* mutations among *Helicobacter pylori* in Japan. Helicobacter 2006;11:243–9.

65. Mégraud F. Current recommendations for Helicobacter pylori therapies in a world of evolving resistance. Gut Microbes 2013;4:541–8.

66. Cuadrado-Lavin A, Salcines-Caviedes JR, Carrascosa MF et al. Antimicrobial susceptibility of *Helicobacter pylori* to six antibiotics currently used in Spain. J Antimicrob Chemother 2012;67:170–3.

67. Wueppenhorst N, Stueger HP, Kist M et al. Identification and molecular characterization of triple- and quadruple-resistant *Helicobacter pylori* clinical isolates in Germany. J Antimicrob Chemother 2009;63:648–53.

68. De Francesco V, Margiotta M, Zullo A et al. Prevalence of primary clarithromycin resistance in *Helicobacter pylori* strains over a 15 year period in Italy. J Antimicrob Chemother 2007;59:783–5.

69. Thung I, Aramin H, Vavinskaya V et al. Review article: the global emergence of *Helicobacter pylori* antibiotic resistance. Aliment Pharmacol Ther 2016;43:514–33.

70. Ierardi E, Giorgio F, Losurdo G et al. How antibiotic resistances could change *Helicobacter pylori* treatment: A matter of geography? World J Gastroenterol 2013;19:8168–80.

71. Hirata K, Suzuki H, Nishizawa T et al. Contribution of efflux pumps to clarithromycin resistance in *Helicobacter pylori*. J Gastroenterol Hepatol 2010;25 Suppl 1:S75–9.

72. Jaka H, Rhee JA, Östlundh L et al. The magnitude of antibiotic resistance to *Helicobacter pylori* in Africa and identified mutations which confer resistance to antibiotics: systematic review and meta-analysis. BMC Infect Dis 2018;18:193.

73. Regnath T, Raecke O, Enninger A et al. Increasing metronidazole and rifampicin resistance of *Helicobacter pylori* isolates obtained from children and adolescents between 2002 and 2015 in southwest Germany. Helicobacter 2017;22.

74. Boyanova L. Prevalence of multidrug-resistant Helicobacter pylori in Bulgaria. J Med Microbiol 2009;58:930–5.

75. Thyagarajan SP, Ray P, Das BK et al. Geographical difference in antimicrobial resistance pattern of *Helicobacter pylori* clinical isolates from Indian patients: *Multicentric study*. J Gastroenterol Hepatol 2003;18:1373–8.

76. van Doorn LJ, Schneeberger PM, Nouhan N et al. Importance of *Helicobacter pylori cagA* and *vacA* status for the efficacy of antibiotic treatment. Gut 2000;46:321–6.

77. Rudi J, Reuther S, Sieg A et al. Relevance of underlying disease and bacterial *vacA* and *cagA* status on the efficacy of *Helicobacter pylori* eradication. Digestion 2002;65:11–5.

78. Sugimoto M, Yamaoka Y. Virulence factor genotypes of Helicobacter pylori affect cure rates of eradication therapy. Arch Immunol Ther Exp 2009;57:45–56.

79. Alarcon-Millan J, Fernandez-Tilapa G, Cortes-Malagon EM et al. Clarithromycin resistance and prevalence of *Helicobacter pylori* virulent genotypes in patients from Southern Mexico with chronic gastritis. Infect Genet Evol 2016;44:190–8.

80. Karabiber H, Selimoglu MA, Otlu B et al. Virulence factors and antibiotic resistance in children with *Helicobacter pylori* gastritis. J Pediatr Gastroenterol Nutr 2014;58:608–12.

81. Rasheed F, Campbell BJ, Alfizah H et al. Analysis of clinical isolates of *Helicobacter pylori* in Pakistan reveals high degrees of pathogenicity and high frequencies of antibiotic resistance. Helicobacter 2014;19:387–99.

82. Sicinschi LA, Correa P, Peek RM et al. CagA C-terminal variations in *Helicobacter pylori* strains from Colombian patients with gastric precancerous lesions. Clin Microbiol Infect 2010;16:369–78.

83. De Francesco V, Margiotta M, Zullo A et al. Claritromycin resistance and *Helicobacter pylori* genotypes in Italy. J Microbiol 2006;44:660–4.

84. Elviss NC, Owen RJ, Xerry J et al. Helicobacter pylori antibiotic resistance patterns and genotypes in adult dyspeptic patients from a regional population in North Wales. J Antimicrob Chemother 2004;54:435–40.

